# An Entropy-based Directed Random Walk for Pathway Activity Inference Using Topological Importance and Gene Interactions

**DOI:** 10.1101/2021.11.05.467449

**Authors:** Tay Xin Hui, Tole Sutikno, Shahreen Kasim, Mohd Farhan Md Fudzee, Shahliza Abd Halim, Rohayanti Hassan, Seah Choon Sen

## Abstract

The integration of microarray technologies and machine learning methods has become popular in predicting pathological condition of diseases and discovering risk genes. The traditional microarray analysis considers pathways as simple gene sets, treating all genes in the pathway identically while ignoring the pathway network’s structure information. This study, however, proposed an entropy-based directed random walk (e-DRW) method to infer pathway activity. This study aims (1) To enhance the gene-weighting method in Directed Random Walk (DRW) by incorporating t-test statistic scores and correlation coefficient values, (2) To implement entropy as a parameter variable for random walking in a biological network, and (3) To apply Entropy Weight Method (EWM) in DRW pathway activity inference. To test the objectives, the gene expression dataset was used as input datasets while the pathway dataset was used as reference datasets to build a directed graph. An equation was proposed to assess the connectivity of nodes in the directed graph via probability values calculated from the Shannon entropy formula. A direct proof of calculation based on the proposed mathematical formula was presented using e-DRW with gene expression data. Based on the results, there was an improvement in terms of sensitivity of prediction and accuracy of cancer classification between e-DRW and conventional DRW. The within-dataset experiments indicated that our novel method demonstrated robust and superior performance in terms of accuracy and number of predicted risk-active pathways compared to the other DRW methods. In conclusion, the results revealed that e-DRW not only improved prediction performance, but also effectively extracted topologically important pathways and genes that are specifically related to the corresponding cancer types.

## Introduction

The accurate prediction of prognosis and metastatic potential of cancer is a major challenge in clinical cancer research. Through the evolution of high-throughput technologies, deoxyribonucleic acid (DNA) microarray analysis can classify tumour samples overriding the traditional diagnostic methods. This technology allows the extraction of a huge amount of molecular information which aids in the discovery of tumour-specific biomarkers. However, the reproducibility of individual gene biomarkers has been challenging as the identified gene markers in one dataset failed to predict the same disease phenotype obtained in other datasets [1]. This discrepancy is usually due to the cellular heterogeneity within tissues, the inherent genetic heterogeneity across patients, and the measurement error in microarray platforms [2]. Besides that, microarray analysis of gene expression data generally produces plenty of genes from patients with the same diseases, hence, leading to a high dimension small sample size problem. All of these factors often decrease the prediction performance and reproducibility of individual gene biomarkers in independent cohorts of patients.

To address unreliable or inconsistent prediction of gene biomarkers in datasets, biological pathway data was introduced to identify robust pathway biomarkers in functional categories [3–6]. As gene products are known to function coordinately in functional modules, the mutual interest between pathway data and gene expression data can extract function-related genes to produce consistent and reproducible biomarkers [7]. Such biomarkers at the functional level can reduce the impact of noise in microarray data by al-lowing a more accurate biological interpretation of the disease-canonical pathway correlations [2]. Generally, network-based microarray analysis can be classified into protein-protein interaction-based (PPI) and pathway-based methods. Both approaches consist of three main steps namely (i) search and sort potential subnetworks or pathways by their discriminative score, (ii) select feature subnetworks or pathways, and (iii) construct a classifier based on the activity of the selected subnetworks or pathways [1]. Both approaches can be distinguished based on the interpretation of the pathway activity.

Multiple studies which incorporated PPI-based methods proposed to identify more robust biomarkers at functional category levels rather than individual genes. For instance, paper [8] proposed a method for determining subnetwork markers based on mutual information or t-scores that measure the relationship between the marker’s activity and class label. Meanwhile, paper [5] applied dynamic programming to determine the top discriminative linear paths and inferred the activity of the subnetwork by gauging normalised log-likelihood ratios (LLRs) of its member genes. Pathway activity scores in pathway-based methods can be calculated based on the t-test statistic score for member genes. Paper [3] employed the mean or median expression value of the member genes to infer the pathway activity. While paper [9] and paper [10] used the first principal component of the expression profile of member genes to evaluate the activity of a given pathway. Conversely, paper [4] proposed pathway activity inference using only a subset of genes in the pathway, called the condition responsive genes (CORGs), in which the combined expression levels can accurately discriminate the phenotypes of interest. Whereas, paper [2] proposed a directed random walk (DRW) to mine the topological importance of genes in a pathway network.

In cancer classification, prior methods that produced significant progress based on the activity of pathways or subnetworks consider pathways as simple gene sets that were treated identically and ignored the structure information of the pathway network [2]. Besides that, the activities of the pathway or subnetworks not only ignored the interactions between the two closest correlated neighbour genes but also failed to reflect on the amount of embodied information based on the expression values of the member genes at different conditions.

To combat the aforementioned issue, this study proposed the entropy-based Directed Random Walk (e-DRW) method to quantify pathway activity using both gene interactions and information indicators based on the probability theory. Apart from mining the topological information of disease genes in a biological network, this method can also reveal the amount of information a gene variable holds in different conditions and infer pathway activity using both gene interactions and entropy probability values. The merged directed pathway network utilises e-DRW to evaluate the topological importance of each gene. Moreover, the expression value of the member genes are inferred based on the t-test statistics scores and correlation coefficient values, whereas, the entropy weight method (EWM) calculates the activity score of each pathway.

The e-DRW method was implemented on the lung cancer dataset (GSE10072) in this study, where the reproducibility of the pathway activities was enhanced and higher ac-curacy was produced. Within-dataset experiments also demonstrated that the e-DRW method was more reliable and robust in predicting clinical outcomes and guiding therapeutic selection. Section 2 of this study presents the datasets and data preprocessing methods used in this experiment. Section 3 presents the research methodology for the proposed approach. The results and detailed discussion of cancer prediction and cancer classification are provided in sections 4 and 5, respectively. Section 6 concludes the study.

## Materials and Methods

The gene expression data is referred to as the input dataset, while the pathway data obtained from public databases are referred to as the reference datasets. This section also briefly describes data preprocessing.

### Gene Expression Data

The GSE10072 for Lung Adenocarcinoma was the gene expression dataset utilised in this analysis [11]. It was downloaded from the National Centre for Biotechnology In-formation (NCBI) Gene Expression Omnibus (GEO) database based on GEO platform GPL96 [12]. It contained a total of 107 samples (Samples number: GSM254625 to GSM254731), where 58 samples represent cancer samples and 49 samples represent normal samples.

### Data Preprocessing

The raw gene expression data consisted of 22,283 genes in rows and 107 samples in columns. The two phases involved in data preprocessing were (i) data cleaning and imputation, and (ii) normalisation of gene expression data. In the first phase, the unwanted and empty values of attributes were removed. Then, rows with incomplete values of at-tributes were imputed with mean values to resolve inconsistencies in data. A total of 1,209 missing values were determined in the GSE10072 lung cancer dataset. The completed dataset following the application of mean imputation was used for inference. However, the rearrangement of data was run through before proceeding to the next phase. The normalisation step in the second phase typically included thresholds or flooring to remove poorly detected probes and log2 transformation to normalise the distribution of probes across the intensity range of the experiment. Whereas, Gene Pattern was used for dataset preprocessing to remove platform noise and genes that have little variation [13]. Following preprocessing, the cleaned dataset contained 12986 genes.

### Directed Pathway Network

The directed pathway network was constructed based on the pathway information obtained from Kyoto Encyclopedia of Genes and Genomes (KEGG) Databases [14]. Firstly, each KEGG pathway was converted into a directed graph using the NetPathMiner software package [15]. A total of 328 human pathways were merged into a directed pathway graph, covering 6,667 nodes and 116,773 directed edges. Each node in the graph represented a gene, while each directed edge represented how genes interacted and controlled each other. The direction of the edge was determined by the type of interaction between two genes found in the KEGG pathway database. For instance, if gene A activates (inhibits) gene B, then A points to B, because one gene influences other genes [16]. This concept is similar to the web page ranking algorithm whereby a web page is important if other pages point to it. Thus, the direction of all edges on the directed pathway graph is reversed to model the conception.

### Methodology

This section describes the approaches in constructing the e-DRW. The conventional DRW method proposed by paper [2] was assessed in this study. Paper [2] used a t-test statistics score as the gene-weighting method to run the algorithm. This study proposed the combination of correlation and t-test values as the gene-weighting method to run e-DRW. Besides that, DRW employed initial probability as a parameter variable to calculate the distribution values for each gene. Entropy was proposed as the weight parameter to reflect the degree of randomness underlying the probability distribution of the directed graph. The application of EWM improved the pathway activity inference method to enhance the sensitivity of cancer prediction and the accuracy of cancer classification. Figure 1 illustrates the workflow of e-DRW to infer pathway activity.

**Figure 1.**
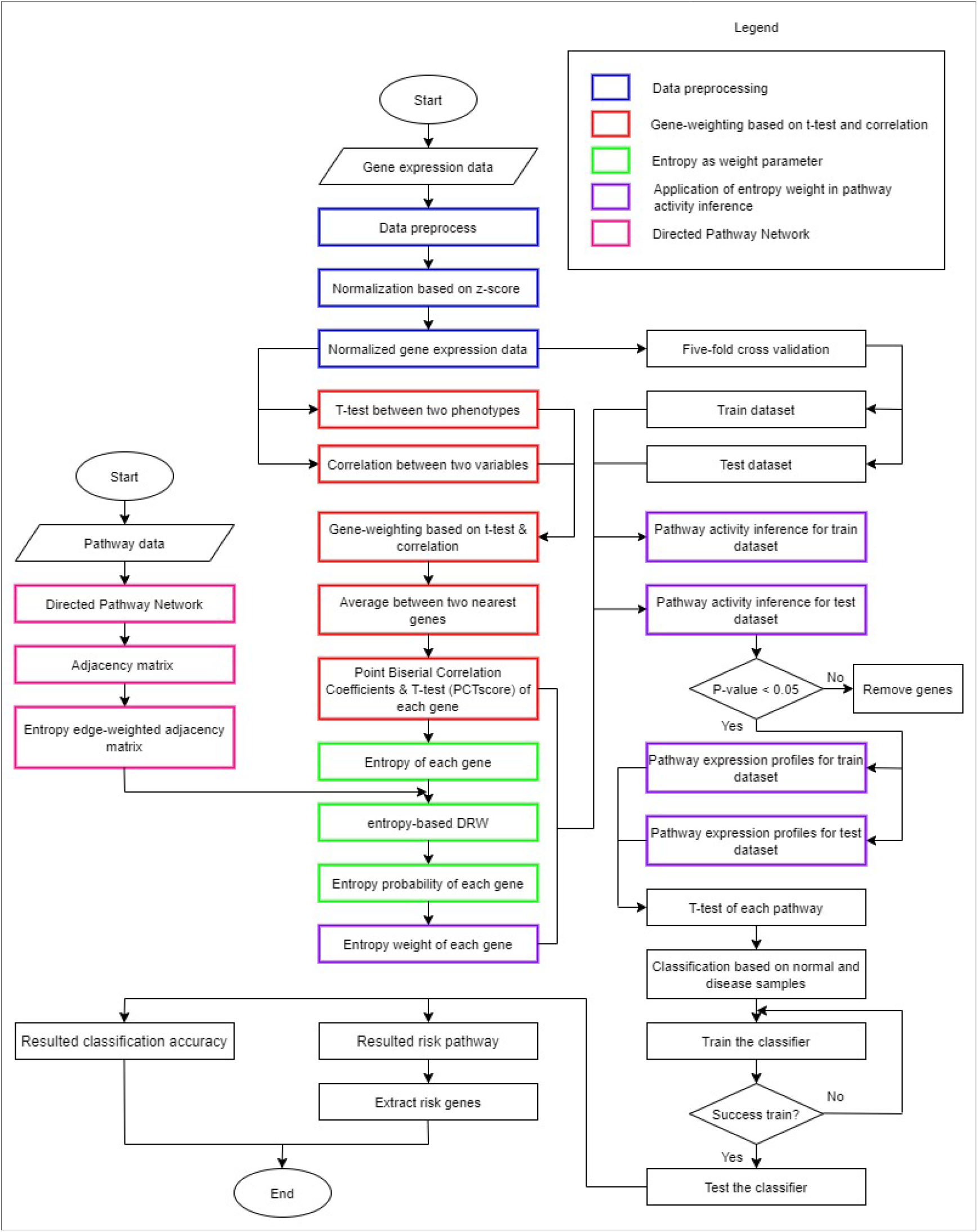
Workflow of e-DRW.

### Gene-weighting Based on Correlation and T-test

This analysis employed a combination of point biserial correlation coefficient and t-test statistic scores as a gene-weighting method. The weighted expressions of the member genes reflected three factors: (1) degree of differential expression of genes between the normal and cancer group; (2) the correlation between a gene expression and class label (normal, cancer); and (3) the average expression values between two closest-correlated neighbour genes. Based on these considerations, a new robust gene-weighting method was proposed in this study.

The normalised expression values of gene, *gi*, in sample *k* was defined as:

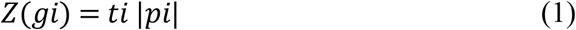

where *ti* is the t-score of gene *gi* calculated using a two-tailed t-test between two phenotypes, while *ρi* is the absolute point biserial correlation coefficient between gene *gi* and class label. *Z(gi)* represents the weighted normalised expression of gene *gi* in sample *k* reflecting the differential expression degree of gene *gi* and its correlation with the phenotype. Larger expression values *(Z(gi))* can be related to higher differential expression and a larger correlation with the phenotype. By employing the averaging method between two nearest neighbour genes proposed by [17], the calculated expression values of gene *gi* were transformed by averaging the gene pair of *gi* and *gj* between two nearest neighbour genes in a directed pathway network as follows:

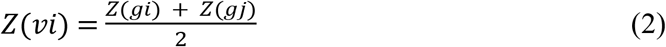

where *Z(gi)* represents the normalised expression values of gene *gi*, *Z(gj)* represents the normalised expression values of gene *gj*, and *Z(vi)* represents the average normalised expression of gene *vi* between two nearest neighbour gene pairs of *gi* and *gj*. The e-DRW was applied for experimentation following the gene-weighting transformation.

### e-DRW

The e-DRW is an improved DRW method that utilises Shannon entropy as an information indicator to calculate the distribution of each node in the directed graph. The amount of information or uncertainty of a sequence in genomic data is calculated using Shannon entropy [18]. Entropy was implemented as a weight parameter in this approach to estimate the variability in expression for a single gene. Node entropy [19] was defined as:

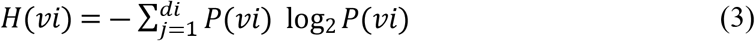

where *P(vi)* represents the probability of the gene. While *vi* was calculated using equation (4):

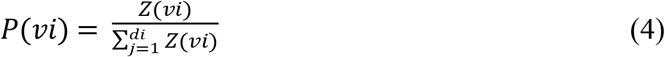

where *Z(vi)* represents the average normalised expression of gene *vi*. In e-DRW, a random walker begins from a single node and transits from its current node either to another randomly selected neighbour (forward) node based on the edge weights or returns to the previous node with probability *r*. According to [2] *r* was set to 0.7. Hence, e-DRW was defined as:

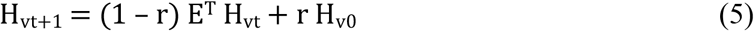

where *Hv_t_* represents the transition probability of ith node transmitted from the *i-1* node. *Hv_0_* is the initial entropy probability vector, *E^T^* is an entropy edge-weighted adjacency matrix developed from the directed graph (with edges), and *Hv_t+1_* denotes the final entropy probability vector.

### Direct Proof of e-DRW

A direct proof of the e-DRW method was determined by calculating the distribution value of the biological pathway, through Leukocyte Transendothelial Migration as demonstrated in a previous study (Sean et al., 2017). The directed biological pathway consists of vertex V = {EPAC → Rap1 → ITGAL → Pyk2 →Vav → RhoA}. Assume a simplified representation of vertex V = {1 → 2 → 3 → 4 → 5 → 6}. Table 1 denotes the weight of nodes after data preprocessing.

**Table 1.** Weight of each node in biological pathway.

| Nodes | Title 2 |
| --- | --- |
| 1 | 2.338914 |
| 2 | 8.47301 |
| 3 | 6.1441 |
| 4 | 3.102989 |
| 5 | 11.388365 |
| 6 | 5.149393 |

The entropy values for each node were estimated before the vector of e-DRW was calculated. Table 2 lists the node entropy based on the probability of gene weight.

**Table 2.** Calculations of node entropy.

| Nodes | Weight | Average Gene Weight, $Z(v_i)$ | Probability of Gene Weight, $P(v_i)$ | Node entropy, $H(v_i)$ |
| --- | --- | --- | --- | --- |
| 1 | 2.338914 | 5.405962 | 0.142272 | 0.400211 |
| 2 | 8.47301 | 7.308555 | 0.192344 | 0.457433 |
| 3 | 6.1441 | 4.623545 | 0.121681 | 0.369789 |
| 4 | 3.102989 | 7.243320 | 0.190627 | 0.455789 |
| 5 | 11.38365 | 8.266522 | 0.217556 | 0.478732 |
| 6 | 5.149393 | 5.149393 | 0.135520 | 0.390758 |
| **Total** |  | 37.997297 | 1 | 2.552712 |

According to Table 2, the average gene weight of the last node (node 6) remained the same as there was no adjacent node connected to node 6 along the pathway. The calculated node entropy was applied in equation (5) to obtain the vector of e-DRW. In the calculation of e-DRW, *r* (restart probability) was set to 0.7, *E^T^* (adjacency matrix) was set to 1, and the initial entropy probability vector was set to 0 as indicated below:

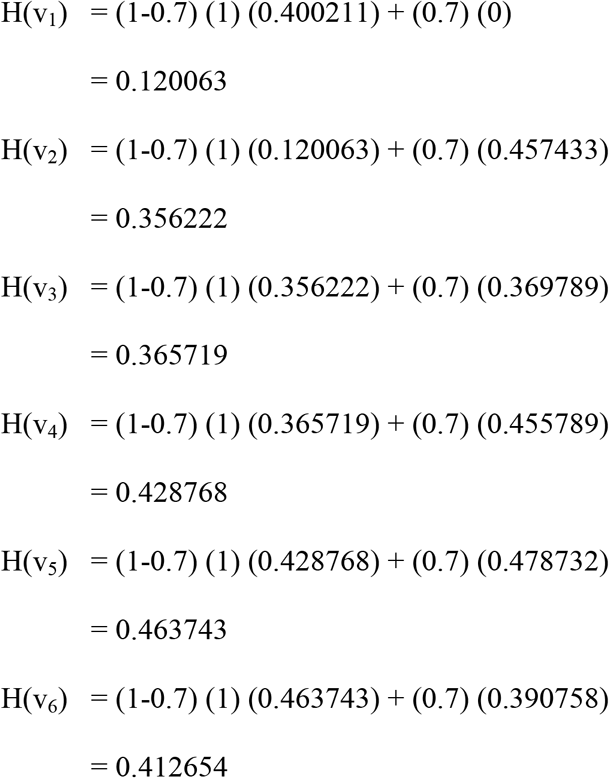

Based on the calculations, the increasing entropy probability vector for random walking proved the feasibility of e-DRW in controlling the randomness of genes according to the biological rules exhibited in Shannon’s entropy of information theory. The increases in entropy of gene *vi* from node 1 to node 6 indicated that it can be used to identify genes, signalling pathways, and novel gene modules within the signalling pathways [20].

### Application of Entropy Weight in Pathway Activity Inference

Paper [2] proposed a pathway activity inference method to infer reproducible pathway activities and robust disease classification. This study, on the other hand, applied EWM in Liu et al.’s proposed inference method to infer the activity score for each pathway. Firstly, the pathway activities were inferred with a set of statistically significant genes (*P* < 0.05). The pathway activity, *a(Pj)*, for training and testing datasets of pathway *Pj* containing *nj* statistically differential expressed genes *{g1, g2, …, gnj*} was calculated as:

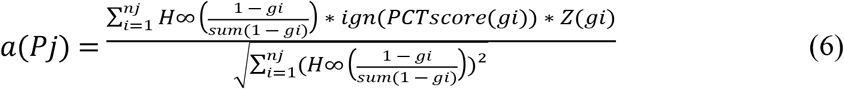

where *H ∞ (gi)* is the entropy weight of gene gi, *sign(PCTscore(gi))* is the sign function of the product of correlation coefficients between gene *gi* and class label, and t-test statistics of gene *gi* from a two-tailed t-test on expression values between two phenotypes, that returns −1 for negative numbers and +1 for positive numbers. Meanwhile, *Z(gi)* is the normalised expression value vector of gene *gi* across the samples in the dataset.

### Classification Evaluation

Classification evaluation was implemented for a within-dataset experiment for the GSE10072 lung cancer dataset. Firstly, the dataset was randomly divided into 5 sets where four-fifths of the samples were used as the training set, while the remaining one-fifth was used as the test set (5-fold cross-validation). Next, to build the classifier, the t-test statistics of the pathway activities on the training dataset was calculated to rank the pathways based on their *P*-values in increasing order. The top 50 pathways were used as candidate features to build the Naïve Bayes model. We constructed the classifier with the pathway that was ranked first. Subsequently, pathways were added sequentially to train the Naïve Bayes model. The performance of the classifier was measured by evaluating its area under the receiver operating characteristics curve (AUC). The added pathway marker was maintained in the feature set if the AUC increased, but was removed if otherwise. This process was repeated for the top 50 pathway markers to optimise the classifier and to yield the best feature set. The performance of the optimised classifier was evaluated on the test set using pathway markers from the best feature set. Each training subset was used sequentially as a feature selection dataset to optimise the classifier. This process was repeated 50 times to ensure unbiased evaluation and to estimate the variation of the AUC. As the final step, the mean AUC across 50 classifiers was estimated to represent the overall performance of the classification method.

## Results

This section presents the classification performance within-dataset experiment. We implemented two pathway-based classification methods, namely, the DRW [2] and Significant Directed Walk (SDW) [17] for comparison purposes. Both the DRW and SDW methodologies were employed in the experimental settings. The Naïve Bayes model was used to evaluate the performance of e-DRW, DRW, and SDW methods. Five-fold cross-validation was also conducted for the three methods, in addition to the estimation of the mean area under the receiver AUC over 50 experiments.

### Classification Performance Within-dataset Experiment

The classification experiments demonstrated that the e-DRW pathway activities yielded reliable predictive accuracy. This method achieved an overall high accuracy across 50 experiments. The average AUC for the GSE10072 lung cancer dataset recorded was approximately 0.995327. Such a consistent performance of e-DRW for within-dataset experiments postulated that the e-DRW pathway activities were more reliable and sensitive to different cohorts of patients and microarray platforms in predicting clinical outcomes in practice. Figure 2 illustrates the classification performance of the e-DRW within-dataset experiment.

**Figure 2.**
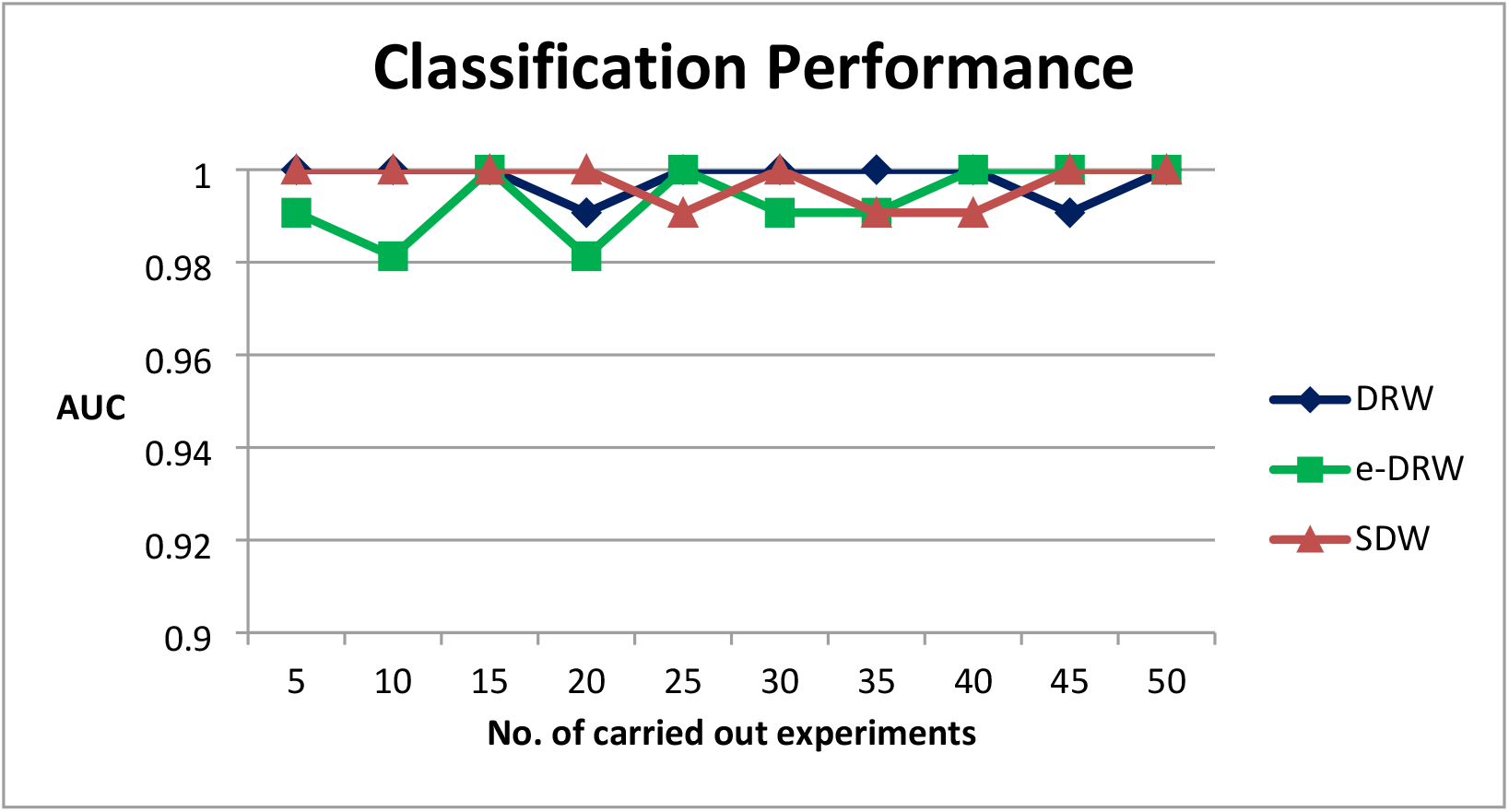
Classification performance of within-dataset experiment.

On the other hand, we also compared classification performance (average AUC) for the GSE10072 lung cancer dataset. The average AUC recorded for DRW and SDW were approximately 0.996262 and 0.994953, respectively. Meanwhile, e-DRW recorded an average AUC value of 0.995327 which fell well between the values of DRW and SDW methods. Figure 3 compares the classification accuracy within-dataset experiments for the three methods.

**Figure 3.**
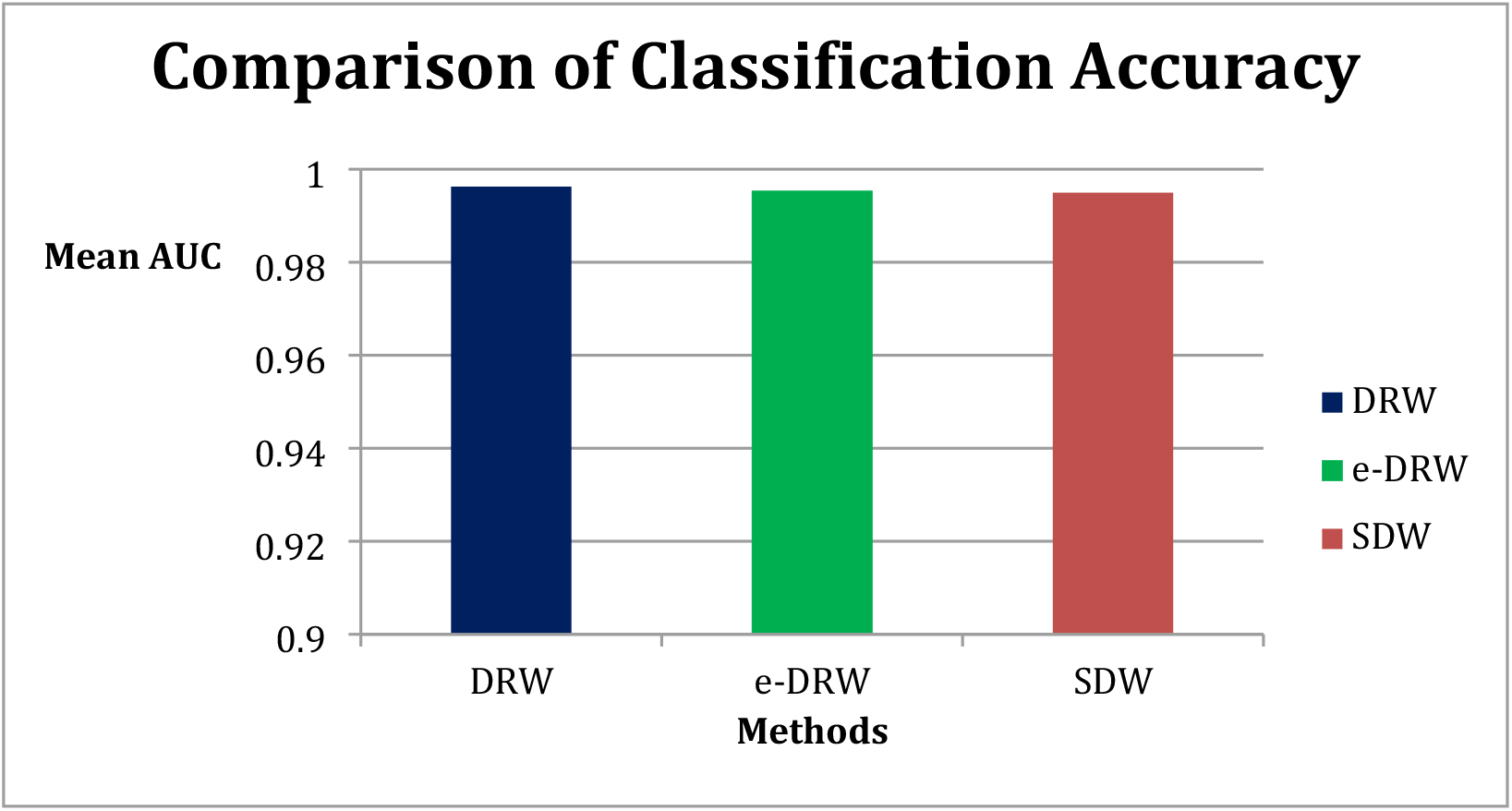
Comparison of classification accuracy.

### Robustness of Risk-active Pathways

The detection of robust risk-active pathways is important in cancer studies. The proposed e-DRW method predicted a total of 23 pathways across 50 experiments. Whereas, the DRW and SDW methods predicted a total of 16 and 19 pathways, respectively, in the lung cancer pathways. Table 3 lists the predicted cancer-related pathways with their respective pathway ID that are involved in various biological processes across the three methods.

**Table 3.** Cancer-related pathway predicted by three methods.

| Methods | No. | Pathway Name | Pathway ID (hsa) | Total Pathways |
| --- | --- | --- | --- | --- |
| e-DRW | 1 | PPAR signalling pathway | 03320 | 23 |
|  | 2 | MAPK signalling pathway | 04010 |  |
|  | 3 | Ras signalling pathway | 04014 |  |
|  | 4 | Rap1 signalling pathway | 04015 |  |
|  | 5 | Calcium signalling pathway | 04020 |  |
|  | 6 | cGMP-PKG signalling pathway | 04022 |  |
|  | 7 | cAMP signalling pathway | 04024 |  |
|  | 8 | Cytokine-cytokine receptor interaction | 04060 |  |
|  | 9 | Chemokine signalling pathway | 04062 |  |
|  | 10 | HIF-1 signalling pathway | 04066 |  |
|  | 11 | Phosphatidylinositol signalling system | 04070 |  |
|  | 12 | Neuroactive ligand-receptor interaction | 04080 |  |
|  | 13 | Autophagy | 04140 |  |
|  | 14 | PI3K-Akt signalling pathway | 04151 |  |
|  | 15 | Adrenergic signalling in cardiomyocytes | 04261 |  |
|  | 16 | Vascular smooth muscle contraction | 04270 |  |
|  | 17 | Axon guidance | 04360 |  |
|  | 18 | Focal adhesion | 04510 |  |
|  | 19 | ECM-receptor interaction | 04512 |  |
|  | 20 | NOD-like receptor signalling pathway | 04621 |  |
|  | 21 | Regulation of actin cytoskeleton | 04810 |  |
|  | 22 | Thyroid hormone signalling pathway | 04919 |  |
|  | 23 | Adipocytokine signalling pathway | 04920 |  |
| SDW | 1 | PPAR signalling pathway | 03320 | 19 |
|  | 2 | MAPK signalling pathway | 04010 |  |
|  | 3 | Ras signalling pathway | 04014 |  |
|  | 4 | Rap1 signalling pathway | 04015 |  |
|  | 5 | Calcium signalling pathway | 04020 |  |
|  | 6 | cGMP-PKG signalling pathway | 04022 |  |
|  | 7 | cAMP signalling pathway | 04024 |  |
|  | 8 | Cytokine-cytokine receptor interaction | 04060 |  |
|  | 9 | Phosphatidylinositol signalling system | 04070 |  |
|  | 10 | Phospholipase D signalling pathway | 04072 |  |
|  | 11 | Neuroactive ligand-receptor interaction | 04080 |  |
|  | 12 | PI3K-Akt signalling pathway | 04151 |  |
|  | 13 | Adrenergic signalling in cardiomyocytes | 04261 |  |
|  | 14 | Vascular smooth muscle contraction | 04270 |  |
|  | 15 | Focal adhesion | 04510 |  |
|  | 16 | Adherens junction | 04520 |  |
|  | 17 | Tight junction | 04530 |  |
|  | 18 | Leukocyte transendothelial migration | 04670 |  |
|  | 19 | Aldosterone synthesis and secretion | 04925 |  |
| DRW | 1 | PPAR signalling pathway | 03320 | 16 |
|  | 2 | MAPK signalling pathway | 04010 |  |
|  | 3 | Ras signalling pathway | 04014 |  |
|  | 4 | Rap1 signalling pathway | 04015 |  |
|  | 5 | Calcium signalling pathway | 04020 |  |
|  | 6 | cGMP-PKG signalling pathway | 04022 |  |
|  | 7 | cAMP signalling pathway | 04024 |  |
|  | 8 | Cytokine-cytokine receptor interaction | 04060 |  |
|  | 9 | Neuroactive ligand-receptor interaction | 04080 |  |
|  | 10 | PI3K-Akt signalling pathway | 04151 |  |
|  | 11 | AMPK signalling pathway | 04152 |  |
|  | 12 | Vascular smooth muscle contraction | 04270 |  |
|  | 13 | Apelin signalling pathway | 04371 |  |
|  | 14 | ECM-receptor interaction | 04512 |  |
|  | 15 | Adherens junction | 04520 |  |
|  | 16 | Human papillomavirus infection | 05165 |  |

Among the three methods, e-DRW predicted the highest number of risk-active pathways that were highly related to the development of various cancers. For instance, the regulation of actin cytoskeleton (hsa04810) is involved in many cancers [2, 21–23]. Besides, the MAPK signalling pathway (hsa04010) and calcium signalling pathway (hsa04020) were known cancer pathways reported in multiple studies [1–2]. Deregulated calcium and PI3K-Akt signalling pathways in many tumours are regarded as a sensitive therapeutic target in several types of cancers including melanoma, lung cancer, and prostate cancer [2, 24]. All the risk-active pathways listed in the table above could assist in cancer detection and in providing new insights for cancer treatment. Based on the number of cancer-related pathways detected, e-DRW was identified to be the most feasible method in detecting cancer modules for cancer classification using gene expression datasets.

## Discussion

This study proposed an entropy-based pathway activity inference scheme for identifying reproducible pathway biomarkers for clinical cancer applications. Previous literature revealed that individual gene markers are less reliable compared to pathway markers, thus, are unable to effectively capture the biological interpretation of gene expression at functional categories [3–6]. Based on the comparative assessment, e-DRW pathway activities were more discriminative and consistent in identifying reproducible pathway biomarkers based on the application of entropy weight. Moreover, EWM is also useful in calculating the amount of information or uncertainty of a biological sequence.

Entropy was implemented as a weight parameter in e-DRW to calculate the distribution values along a biological pathway. This weight parameter is important in revealing the biological insights for a gene and a pathway. With proven results following implementation, entropy also disclosed the essential biological information behind the concepts of information theory. Increasing entropy vector values proved to be related to cancer modules in signalling pathways [20]. Although different representations of weight variables can influence the calculation of vectors along the pathway, entropy values re-main important in the implementation of e-DRW for cancer classification.

Based on the classification performance, the AUCs of the e-DRW method were significantly higher and stable across the experiments. The reliable performance of the e-DRW pathway activities could be attributed to the strategy of weighting genes according to their topological importance and correlation values between the nearest-neighbour genes. The gene-weighting method based on t-test and correlation can greatly magnify the signals of essential genes whose expression levels may have a large impact on the pathway while weakening the differential signals of genes that only appear downstream or have a minor impact on the system. Therefore, the e-DRW approach could alleviate the noise caused by sample heterogeneity or technical measurements, resulting in more reproducible pathway activities.

Based on the comparison experiments, the average AUC of e-DRW was better in terms of cancer classification due to higher accuracy compared to DRW. Comparatively, e-DRW also recorded more risk-active pathways than that of the 7 pathways reported by DRW. Overall, e-DRW was more effective in gene prediction as it was sensitive compared to DRW.

## Conclusions

In cancer studies, accurate prediction of cancer is crucial for the diagnosis and prognosis of clinical therapy. We proposed an e-DRW approach based on entropy (weight parameter) for cancer classification. The three enhancements based on paper [2]’s work and the proposed e-DRW were proven to be effective in inferring pathway activities and accurate cancer classification. The three proposed enhancements included (1) gene-weighting based on correlation and t-test, (2) entropy as parameter variable, and (3) application of entropy weight in pathway activity inference. Gene-weighting method in e-DRW incorporated t-test statistics scores and correlation coefficient values to weigh each gene in a directed pathway network. This weighting strategy not only reflects the degree of the differential expression of genes between normal and cancer groups but also considers the correlation values between two nearest neighbour genes in a directed graph. Secondly, entropy was introduced as a weight variable to measure the vector along the biological pathway. This metric was useful in disclosing essential biological information behind the concepts of information theory. Next, EWM was utilised in conventional DRW along with the product of t-test and correlation values of each gene to calculate the activity score for each pathway. The EWM was based on the idea that superior weight indicator information was more constructive than that of the lower indicator information [25]. Finally, the five-fold cross-validation was employed to train the classifier and classify the significant pathways detected by e-DRW. In conclusion, the proposed approach was more effective and feasible for cancer classification compared to other DRW methods.

## Acknowledgements

This study was supported by Universiti Tun Hussein Onn Malaysia under REGG FASA 1/2021 (VOT NO. H888). This work also was supported/funded by the Ministry of Higher Education under Fundamental Research Grant Scheme (FRGS/1/2020/ICT02/UTM/03/1).

## Notes

### Competing Interest Statement

The authors have declared no competing interest.

## References

1. Xu, P., Zhao, G., Kou, Z., Fang, G., & Liu, W. (2020). Classification of cancers based on a comprehensive pathway activity inferred by genes and their interactions. IEEE Access, 8, 30515–30521.

2. Liu, W., Li, C., Xu, Y., Yang, H., Yao, Q., Han, J., … & Li, X. (2013). Topologically inferring risk-active pathways toward precise cancer classification by directed random walk. Bioinformatics, 29(17), 2169–2177.

3. Guo, Z. et al. Towards precise classification of cancers based on robust gene functional expression profiles. BMC Bioinformatics 6, 58 (2005).

4. Lee, E., Chuang, H. Y., Kim, J. W., Ideker, T. & Lee, D. Inferring pathway activity toward precise disease classification. PLoS Comput Biol 4, e1000217 (2008).

5. Su, J., Yoon, B. J. & Dougherty, E. R. Accurate and reliable cancer classification based on probabilistic inference of pathway activity. PLoS One 4, e8161 (2009).

6. Kim, S., Kon, M. & DeLisi, C. Pathway-based classification of cancer subtypes. Biol Direct 7, 21 (2012).

7. Y. Wu, H. Huang, Q. Wu, A. Liu, and T. Wang, ‘‘A risk defense method based on microscopic state prediction with partial information observations in social networks,’’ J. Parallel Distrib. Comput., vol. 131, pp. 189–199, Sep. 2019.

8. H. Y. Chuang, E. Lee, Y. T. Liu, D. Lee, and T. Ideker, ‘‘Network-based classification of breast cancer metastasis,’’ Mol. Syst. Biol., vol. 3, no. 1, p. 140, Oct. 2007.

9. Tomfohr, J., Lu, J., & Kepler, T. B. (2005). Pathway level analysis of gene expression using singular value decomposition. BMC bioinformatics, 6(1), 1–11.

10. Bild, A. H., Yao, G., Chang, J. T., Wang, Q., Potti, A., Chasse, D., … & Nevins, J. R. (2006). Oncogenic pathway signatures in human cancers as a guide to targeted therapies. Nature, 439(7074), 353–357.

11. Landi,M.T. et al. (2008) Gene expression signature of cigarette smoking and its role in lung adenocarcinoma development and survival. PLoS One, 3, e1651.

12. R. Edgar, M. Domrachev, and A. E. Lash, ‘‘Gene expression omnibus: NCBI gene expression and hybridization array data repository,’’ Nucleic Acids Res., vol. 30, pp. 207–210, Jan. 2002.

13. Kuehn, H., Liberzon, A., Reich, M., & Mesirov, J. P. (2008). Using GenePattern for gene expression analysis. Current protocols in bioinformatics, 22(1), 7–12.

14. Kanehisa,M. and Goto,S. (2000) KEGG: kyoto encyclopedia of genes and genomes. Nucleic Acids Res., 28, 27–30.

15. Mohamed, A., Hancock, T., Nguyen, C. H., & Mamitsuka, H. (2014). NetPathMiner: R/Bioconductor package for network path mining through gene expression. Bioinformatics, 30(21), 3139–3141.

16. Draghici,S. et al. (2007) A systems biology approach for pathway level analysis. Genome Res., 17, 1537–1545.

17. Seah, C. S., Kasim, S., Fudzee, M. F. M., & Mohamad, M. S. (2017, September). A direct proof of significant directed random walk. In IOP Conference Series: Materials Science and Engineering (Vol. 235, No. 1, p. 012004). IOP Publishing.

18. Akhter, S., Bailey, B. A., Salamon, P., Aziz, R. K., & Edwards, R. A. (2013). Applying Shannon’s information theory to bacterial and phage genomes and metagenomes. Scientific reports, 3(1), 1–7.

19. Chen, Z., Dehmer, M., Emmert-Streib, F., & Shi, Y. (2015). Entropy of Weighted Graphs with Randi c Weights. Entropy, 17(6), 3710–3723.

20. West, J., Bianconi, G., Severini, S., & Teschendorff, A. E. (2012). Differential network entropy reveals cancer system hallmarks. Scientific reports, 2(1), 1–8.

21. Wang,W. et al. (2004) Identification and testing of a gene expression signature of invasive carcinoma cells within primary mammary tumors. Cancer Res., 64, 8585–8594.

22. Chuma,M. et al. (2004) Overexpression of cortactin is involved in motility and metastasis of hepatocellular carcinoma. J. Hepatol., 41, 629–636.

23. Yamaguchi, H., & Condeelis, J. (2007). Regulation of the actin cytoskeleton in cancer cell migration and invasion. Biochimica et Biophysica Acta (BBA)-Molecular Cell Research, 1773(5), 642–652.

24. K.-Q. Liu, Z.-P. Liu, J.-K. Hao, L. Chen, and X.-M. Zhao, ‘‘Identifying dysregulated pathways in cancers from pathway interaction networks,’’ BMC Bioinf., vol. 13, no. 1, p. 126, 2012.

25. Kumar, R., Singh, S., Bilga, P. S., Jatin, K., Singh, J., Singh, S., … & Pruncu, C. I. (2021). Revealing the benefits of entropy weights method for multi-objective optimization in machining operations: A critical review. journal of materials research and technology.

